## Supplementary material for "The chromatin factors SET-26 and HCF-1 oppose the histone deacetylase HDA-1 in longevity and gene regulation in *C. elegans*": Description of Supplementary Data Files

Emerson et al

File Name: Supplementary Data 1

Description: CUT&RUN binding sites (peaks and genes) under basal conditions in *C. elegans*

File Name: Supplementary Data 2

Description: Integration of RNA-seq and CUT&RUN data in *C. elegans* mutants

File Name: Supplementary Data 3

Description: CUT&RUN binding sites (peaks and genes) in mutant backgrounds in *C. elegans*
