## Supplementary Information for "The chromatin factors SET-26 and HCF-1 oppose the histone deacetylase HDA-1 in longevity and gene regulation in *C. elegans*"

Supplementary Information for Emerson et al.

**Supplementary Information**

Pages

Supplementary Figures	2-10

Supplementary Methods	11-13

Supplementary Tables	14-16

Supplementary References	17

**Supplementary Fig. 1: Quality control and whole worm profiles for SET-26 and HCF-1 binding.** **a** Alignment of the HCF-1 binding motif (DHNY) and immediate upstream sequence identified in MLL5 (by Zhou et al.^1^) and the corresponding amino acid sequences from SET-26 and KMT2E showing conservation from *C. elegans* to *H. sapiens*. **b-c** Survival curves for worms carrying epitope-tagged (**b**) *hcf-1* (*hcf-1::gfp::3xflag*) or (**c**) *set-26* (*set-26::ha)* compared to wildtype controls and loss of function mutants of *hcf-1* or *set-26* from one representative experiment. **d** Pearson’s correlation of the CUT&RUN profiles for either SET-26 or HCF-1 binding in the *glp-1(-)* mutant background in two independent replicates. **e-f** Annotation of the proportion of somatic (**e**) SET-26 or (**f**) HCF-1 peak regions that overlap with the indicated genomic features. **g** Pearson’s correlation of the CUT&RUN profiles for SET-26 or HCF-1 binding in the wildtype background containing germlines in two independent replicates. **h-i** Annotation of the proportion of whole worm (**h**) SET-26 or (**i**) HCF-1 peak regions that overlap with the indicated genomic features. **j** Venn diagram showing whole worm SET-26 and HCF-1 peaks in the wildtype background, and 5,280 peaks show overlap by 1bp or more. **k-l** Venn diagrams showing the number of genes associated with either (**k**) SET-26 or (**l**) HCF-1 peaks in somatic (*glp-1(-)* mutant) samples compared to whole worm (wildtype) samples. **m-n** GO term analysis from Wormcat for the (**m**) 1,141 genes bound by SET-26 or (**n**) 1,526 genes bound by HCF-1 that were identified only in wildtype whole worm samples containing germlines and not in *glp-1(-)* mutants. *** indicates *p<1x10^-15^* and overlap is higher than expected by chance, as calculated by hypergeometric test for peak overlap (in **j**) and Fisher’s Test for gene overlap (in **k** and **l**). In (**m-n**), Wormcat *p* values are determined by Fisher test with FDR correction. Quantitative data are provided in Source Data and peaks and gene sets are provided in Supplementary Data 1.

**Supplementary Fig. 2: Quality control for RNA-seq analysis. a** PCA plot showing correlation between RNA-seq replicates for *glp-1(-), glp-1(-);hcf-1(-),* and *glp-1(-);set-26(-)* mutants at day 1 of adulthood. **b-c** Wormcat GO enrichment analysis for genes commonly (**b**) upregulated (485 genes) or (**c**) downregulated (602 genes) in RNA expression in both *glp-1(-);hcf-1(-),* and *glp-1(-);set-26(-)* mutants compared to single *glp-1(-)* mutants as determined in Fig. 2a and b. Wormcat *p* values are determined by Fisher test with FDR correction. Gene sets are provided in Supplementary Data 2.

**Supplementary Fig. 3: HCF-1 requires SET-26 primarily in somatic cells for its recruitment to chromatin. a** Screenshot from IGV shows somatic HCF-1 binding in a portion of Chromosome V, with binding and peak calls either normalized to antibody background (top) as in Fig. 1d-e, or to H3 (bottom) as in Fig. 3a. CUT&RUN profiles were captured by CUT&RUN of *hcf-1::gfp::3xflag* worms in either the *glp-1(-)* mutant background (top) or with *glp-1* RNAi (bottom). **b** Pearson’s correlation of the CUT&RUN profiles for HCF-1 or H3 in controls (*hcf-1::gfp::3xflag* worms) or *set-26(-)* mutants grown on *glp-1* RNAi in two independent replicates. **c** Normalized counts of *hcf-1* RNA expression from RNA-seq of day 1 adult *glp-1*, *glp-1(-);set-26(-)*, or *glp-1(-);hcf-1(-)* mutants as determined by DESeq2. Each dot represents one biological replicate. **d** Representative replicate of immunoblotting experiments targeting HCF-1 or H3 in either day 1 adult controls (*hcf-1::gfp::3xflag* worms) or *set-26(-)* mutants in wildtype or *glp-1(-)* mutant backgrounds. N2 and *glp-1(-)* untagged worms were used as negative controls for the FLAG antibody (lanes 1 and 4). **e** Quantification of immunoblotting replicates comparing HCF-1 levels normalized to H3 in *set-26(-)* mutants versus controls in wildtype (whole worm) or germline-less *glp-1(-)* mutant (somatic) backgrounds. N = 2 independent experiments. **f** Pearson’s correlation of the CUT&RUN profiles for HCF-1 or H3 in *glp-1(-)* single mutants or *glp-1(-);set-26(-)* double mutants in two independent replicates. **g** Metaplot (top) and heatmap (bottom) of HCF-1 signal in somatic HCF-1 binding sites and surrounding 2kb up-and downstream in either *glp-1(-)* single mutants or *glp-1(-);set-26(-)* double mutants. **h** Volcano plot of HCF-1 binding regions determined by DiffBind to be significantly different (pink, FDR <= 0.05) or unchanged (blue, FDR >0.05) in *glp-1(-);set-26(-)* mutants compared to *glp-1(-)* single mutants. **i** Pearson’s correlation of the CUT&RUN profiles for HCF-1 or H3 control in either wildtype worms or *set-26(-)* mutants grown on empty vector (E.V) RNAi in two independent replicates. **j** Metaplot (top) and heatmap (bottom) of HCF-1 signal in whole worm HCF-1 binding sites and surrounding 2kb up-and downstream in either WT or *set-26(-)* mutants grown on E.V. control RNAi. **k** Volcano plot of HCF-1 binding regions determined by DiffBind to be significantly different (pink, FDR <= 0.05) or unchanged (blue, FDR >0.05) in *set-26(-)* mutants compared to wildtype worms grown on E.V. control RNAi. In (**c**), *** indicates FDR <0.0005 and *n.s.* FDR >0.05 as determined by DESeq2. In (**e**), *n.s.* represents p>0.05 as determined by two-tailed t-test and error bars represent standard errors. In (**h**) and (**k**), DiffBind FDR values are calculated using DESeq2. Quantitative data are provided in Source Data. Gene sets and differential peaks are provided in Supplementary Data 3.

**Supplementary Fig. 4: The requirement for HCF-1 in a subset of SET-26 binding sites primarily occurs in somatic cells.** **a-b** Pearson’s correlation of CUT&RUN profiles for SET-26 or H3 in controls (*set-26::ha* worms) or *hcf-1(-)* mutants grown on (**a**) *glp-1* RNAi or (**b**) E.V. control RNAi in two independent replicates. **c** Metaplot (top) and heatmap (bottom) of SET-26 signal in whole worm SET-26 binding sites and surrounding 2kb up-and downstream in either controls or *hcf-1(-)* mutants grown on E.V. control RNAi. **d** Volcano plot of SET-26 binding regions determined by DiffBind to be significantly different (pink, FDR <= 0.05) or unchanged (blue, FDR >0.05) in *hcf-1(-)* mutants compared to controls grown on E.V. control RNAi. DiffBind FDR values are calculated using DESeq2. **e-f** Wormcat GO enrichment analysis for the (**e**) 122 genes identified in Fig. 4h or (**f**) 79 genes identified in Fig. 4i that are upregulated or downregulated respectively in germline-less *set-26(-)* and *hcf-1(-)* mutants, and which display decreased somatic SET-26 and HCF-1 binding in the opposite mutant. Wormcat *p* values are determined by Fisher test with FDR correction. Gene sets and differential peaks are provided in Supplementary Data 2 and 3.

**Supplementary Fig. 5: Validation of *hda-1* requirement for *set-26(-)* and *hcf-1(-)* mutant longevity and HDA-1 binding profiles.** **a-b** *hda-1* RNA expression measured by qRT-PCR in day 3 adults treated with (**a**) *hda-1* RNAi from the Ahringer library or (**b**) newly constructed *hda-1 3’ UTR* RNAi initiated on day 1 of adulthood in WT control worms, *set-26(-)* mutants, or *hcf-1(-)* mutants. Fold change represents the change in *hda-1* RNA levels in worms on *hda-1* RNAi relative to each genotype grown on E.V. RNAi. **c** Survival curves for wildtype controls, *set-26(-)*, and *hcf-1(-)* mutants on E.V. control RNAi and *hda-1* *3’ UTR* RNAi from one representative experiment. *hda-1* RNAi was initiated on day 1 of adulthood. **d** Pearson’s correlation of the CUT&RUN profiles for HDA-1 (*hda-1::gfp::ha* worms) in a *glp-1(-)* mutant background in two independent replicates. **e** Annotation of the proportion of somatic HDA-1 peak regions that overlap with the indicated genomic features. **f** Pearson’s correlation of the whole-worm CUT&RUN profiles for HDA-1 (*hda-1::gfp::ha* worms) in a wildtype background in two independent replicates. **g** Annotation of the proportion of whole worm HDA-1 peak regions that overlap with the indicated genomic features. **h** Venn diagram showing SET-26, HCF-1, and HDA-1 peaks found from whole worm wildtype samples and the 3,323 peaks that overlap by 1bp or more. **i** Venn diagram showing the number of genes identified as bound by HDA-1 in somatic (*glp-1(-)* mutant) samples compared to whole worm (wildtype) samples. In (**a** and **b**) * indicates *p<0.05* in one-tailed t-test. Error bars represent standard errors, N=2 independent experiments for all qRT-PCR experiments. In (**h** and **i**), *** indicates *p<1x10^-15^* and the overlap is higher than expected by chance, as calculated by hypergeometric test for peak overlap (in **h**) and Fisher’s Test for gene overlap (in **i**). Quantitative data are provided in Source Data and gene sets are provided in Supplementary Data 1.

**Supplementary Fig. 6: HDA-1 recruitment to chromatin is more dramatically affected by loss of *set-26* in whole worms than in germline-less worms. a-b** Pearson’s correlation of the CUT&RUN profiles for either HDA-1 or H3 in controls (*hda-1::gfp::ha* worms), *set-26(-)* mutants, or *hcf-1(-)* mutants grown on (**a**) *glp-1* RNAi or (**b**) E.V. control RNAi in two independent replicates. **c** Metaplot (top) and heatmap (bottom) of z-scores representing normalized HDA-1 signal in whole worm HDA-1 binding sites and surrounding 2kb up-and downstream in controls, *set-26(-)* mutants, or *hcf-1(-)* mutants grown on E.V. control RNAi. **d-e** Volcano plots of HDA-1 binding regions determined by DiffBind to be significantly different (pink, FDR <= 0.05) or unchanged (blue, FDR >0.05) in (**d**) *set-26(-)* or (**e**) *hcf-1(-)* mutants compared to controls grown on E.V. control RNAi. Diffbind FDR values in are calculated using DESeq2. Gene sets and differential peaks are provided in Supplementary Data 3.

**Supplementary Fig. 7. HDA-1-dependent gene expression and mitoUPR activation in *set-26(-)* and *hcf-1(-)* mutants. a** Representative replicate of immunoblotting experiments targeting HDA-1 or H3 in day 3 adults in either controls (*hda-1::gfp::ha* worms) or germline-less *glp-1(-)* mutants exposed to control E.V. RNAi or *hda-1* RNAi initiated on day 1 of adulthood. N2 and *glp-1(-)* untagged worms were used as negative controls for the HA antibody (lanes 1 and 4). **b** Quantification of immunoblotting replicates comparing HDA-1 level normalized to H3 in worms exposed to *hda-1* RNAi versus E.V. controls in wildtype (whole worm) or germline-less *glp-1(-)* mutant (somatic) backgrounds. N = 2 independent experiments. **c-d** PCA plot showing correlation between RNA-seq replicates for *glp-1(-), glp-1(-);hcf-1(-),* and *glp-1(-);set-26(-)* mutants aged on E.V. or *hda-1* RNAi initiated on day 1 of adulthood. Samples were collected at either (**c**) day 3 or (**d**) day 12 of adulthood. **e-f** Normalized read counts as determined by DESeq2 of (**e**) *hsp-6* or (**f**) *hsp-60* RNA expression from RNA-seq of day 1 adult *glp-1(-)*, *glp-1(-);set-26(-)*, and *glp-1(-);hcf-1(-)* mutants. Each dot represents one biological replicate. *hsp-6* RNA displays a fold change increase of 5.46 in *set-26(-)* mutants and 3.12 in *hcf-1(-)* mutants respectively, while *hsp-60* RNA displays a fold change increase of 5.82 in *set-26(-)* mutants and 2.79 in *hcf-1(-)* mutants respectively. In (**b**), * represents *p<0.05* as determined by one-tailed t-test and error bars represent standard errors. In (**e** and **f**), * represents *FDR<0.05* and ** represents *FDR<1x10^-5^* as determined by DESeq2. Quantitative data are provided in Source Data.

**Supplementary Methods**

**CUT&RUN data analysis (continued)**

In the linux environment, adaptor sequences were trimmed and low quality reads were filtered out from sequencing files using Trim Galore! (v0.6.5), which utilizes Cutadapt (v3.4)^2^ and FastQC (v0.11.8), with the settings –paired –q 20 –fastqc. Trimmed paired-end sequencing files were then aligned to the ce11/WBcel235 *C. elegans* reference genome using bowtie2 (v2.4.3)^3^ with the options –very-sensitive-local –no-unal –no-mixed –no-discordant –phred33 -p 4 -I 10 -X 700. The resulting sam files were then converted to bam files using ‘samtools view’ command in SAMTools (v1.14)^4^ with the options -hSb -F 4. Since keeping duplicated reads is recommended for CUT&RUN, they were not removed^5,6^. The individual bam files corresponding to two biological replicates were considered both individually (for correlation analysis and DiffBind) and combined (for heatmaps, metaplots, and visualization on IGV (v2.4.19)^7^) using the ‘samtools merge’ command in SAMTools for downstream analysis. The bam files were then sorted using ‘samtools sort’ command and indexed using the ‘samtools index’ command in SAMTools using default settings.

Indexed bam files were used for narrow peak calling with MACS2 (v2.1.4)^8^ using the settings -f BAM - g ce –call-summits –keep-dup all -q 0.01 -m 5 50 –nomodel. When calling peaks for factor binding to characterize binding sites (as in Fig. 1), peaks of each factor were called using combined replicates of factor binding against a control bam file of two combined biological replicates of experiments targeting the tag of interest in the N2 background (i.e. CUT&RUN with an anti-FLAG or anti-HA antibody in N2 or *glp-1(-)*, which produces background antibody binding tracks). When calling peaks for factor binding to directly compare the degree of binding in two genotypes (as in Fig. 3, 4, and 6), peaks were called with a control file representing H3 binding in each genotype (i.e. *hcf-1::gfp::3xflag* *set-26(-)* strain subjected to H3 CUT&RUN in parallel to FLAG). This process was repeated with the individual bam files for each replicate and the merged bam files representing both replicates as discussed above. The number of peaks discussed represents the number of unique peaks called (after removing duplicate locations with multiple summits called by MACS2) and results from peak calling using merged bam files.

To generate bigwig files for visualization, combined bam files from both biological replicates were put through the ‘bamCompare’ command in deepTools (v3.3)^9^ with the settings –binSize 20 –operation log2 –scaleFactors Method None –normalizeUsing CPM –numberOfProcessors 8 –outFileFormat bigwig –smoothLength 60 –extendReads –centerReads, and with –bamfile2 representing a bam file with combined replicates of CUT&RUN experiments targeting either the tagged factor of interest in untagged control worms (N2 or *glp-1(-)*) or H3, as indicated in figure legends.

To determine the correlation between biological replicates, the mapped read count in 2 kb sliding windows was obtained for each bam file corresponding to a single replicate using the command ‘bedtools multicov’ in BEDtools (v2.29.2)^10^ using a reference of the ce11/WBcel235 genome broken into 2 kb windows generated with the ‘bedtools makewindows’ command. The readcount of each sequencing file in each 2 kb window was uploaded into RStudio, converted into log10, and then the Pearson’s correlation was calculated between each replicate using the ‘cor’ function in R. The correlations were saved in a matrix and plotted using the ‘heatmap.2’ function within the gplots package (v3.1.3) in R.

To find overlapping peaks, peak regions called by MACS2 for factor binding vs antibody background were uploaded into the R environment in the package ChIPpeakAnno (v3.28.1)^11^ and used as input for the command ‘findOverlapsOfPeaks’ with the settings ‘minoverlap=1, connectedPeaks=”keepAll”). The overlapping peaks were then plotted using the ‘makeVennDiagram’ command, and a hypergeometric test was performed to test the significance of overlap between the peaksets using the option ‘totalTest=30000’.

To annotate the genomic features associated with peaks, the same peaks were used as input for the ‘assignChromosomeRegion’ command in ChIPpeakAnno using the default settings and the annotation data of TxDb.Celegans.UCSC.ce11.ensGene downloaded into the R environment from Bioconductor (v3.15). To produce density plots of factor binding around the TSS, the same peak regions were used as input for the ‘binOverFeature’ command in ChIPpeakAnno, using the default settings with radius = 3000 and the annotation data of TxDb.Celegans.UCSC.ce11.ensGene. To produce density plots of SET-26 and HCF-1 binding within common peak regions, the overlapping peaks identified from the ‘findOverlapsOfPeaks’ command were recentered with a width of 1. The factor binding signal extracted from bigwig files containing combined replicates of normalized factor binding versus antibody background were used as input (cvglists) for the ‘featureAlignedSignal’ command in ChIPpeakAnno, with the recentered overlapping peaks as the reference frame of which to plot (feature.gr).

To annotate CUT&RUN peaks with associated genes, the peaks of each factor binding versus antibody background were uploaded to ChIPpeakAnno and used as input for the ‘annotatePeakInBatch’ function in ChIPpeakAnno, with the settings of ‘output = “nearestBiDirectionalPromoters”, bindingRegion = c(-2000, 500)’ and the annotation data of TxDb.Celegans.UCSC.ce11.ensGene. The process was repeated for regions found to be significantly differentially bound by DiffBind. BioVenn ([www.biovenn.nl](http://www.biovenn.nl)) was used for generating Venn Diagrams^12^, and the significance of overlapping genes were determined by Fisher’s exact test calculated in RStudio. WormCat 2.0 ([www.wormcat.com](http://www.wormcat.com)) was used for gene ontology enrichment analysis, where the p values are determined by Fisher’s test with FDR correction^13^.

To produce z-score normalized bigwig files of factor binding in different genotypes to use as input for heatmaps and metaplots, combined bam files from both biological replicates of factor binding were first converted to bedgraphs using the ‘bamCompare’ command with the settings –binSize 20 –operation log2 –scaleFactorsMethod readCount –numberOfProcessors 8 –outFileFormat bedgraph –smoothLength 60 –extendReads, and with –bamfile2 representing a bam file with combined replicates of CUT&RUN experiments targeting H3 in parallel in the same genotype. The resulting log2 signal bedgraph files were then uploaded into R (v4.1.2) using RStudio (v2022.02.0+443) and converted into z-scores using the ‘scale’ function in R. Z-score converted bedgraphs were uploaded back into the Linux environment, sorted using the command ‘do sort -k1,1 -k2,2n’, and converted to bigwig files using the UCSC bedGraphToBigWig program with default settings.

To generate heatmaps and metaplots, computeMatrix from deepTools was used to generate a standardized matrix of z-score factor binding in predetermined peak regions with the sub-command ‘scale-regions’ to scale all peak regions to the same size for visualization, with settings -b 2000 -a 2000 –binSize 20 –sortRegions keep –regionBodyLength 1000 -p 4 -p max/2. The bed files of predetermined peak regions (--regionsFileName argument) were generated from MACS2 peak calling of factor vs antibody background (e.g. FLAG signal in the *hcf-1::gfp::3xflag* strain vs FLAG signal in N2) in combined replicates as described above. The z-score normalized bigwig files generated above were used as the files containing scores to be plotted (--scoreFileName argument). The resulting matrix files were used as input for the ‘plotHeatmap’ command in deepTools.

To compare differentially bound regions between two genotypes, sorted bam files of factor binding and control immunoprecipitations (against H3) and peak calls from MACS2 of factor binding against H3 controls for each of two biological replicates were read into the R environment and analyzed using the DiffBind package (v3.4.11)^14^ with the comparison: (factor binding in mutant - H3 binding in mutant) - (factor binding in control - H3 binding in control). The standard DiffBind workflow was used, with dba.count settings of ‘minOverlap=2, bUseSummarizeOverlaps = TRUE’, and Volcano plots and p values generated using DBA_DESEQ2 to assess significance. Regions were considered significantly differentially bound if they were assigned an FDR value of less than 0.05 by the DESeq2 algorithm utilized by DiffBind.

**Supplementary Table 1. *C. elegans* strains**

| **Strain Name** | **Genotype** | **Source** |
| --- | --- | --- |
|  | N2 | CGC |
|  | WT | Wild-type sibling of IU687-IU689 (N2 background) |
| IU687.1 | *hcf-1(pk924) set-26(tm2467)* | Generated by genetic cross in this study |
| IU688.1 | *set-26(tm2467)* | Generated by genetic cross in this study |
| IU689.1 | *hcf-1(pk924)* | Generated by genetic cross in this study |
|  | *glp-1(e2141)* | *glp-1* single mutant sibling of IU701.1 and parental strain of IU714.1 |
| IU714.1 | *glp-1(e2141);set-26(tm2467)* | Generated by genetic cross in this study |
| IU701.2 | *glp-1(e2141);hcf-1(pk924)* | Generated by genetic cross in this study |
| IU716.1 | *rwIs33*[*hcf-1*::*gfp*::3x*flag*] | C-terminal CRISPR endogenous knock-in generated in this study |
| IU717.1 | *rwIs27*[*set-26*::*ha*] | C-terminal CRISPR endogenous knock-in generated in this study |
| IU723.1 | *syb5185*[*hda-1::gfp::ha*] | C-terminal CRISPR endogenous knock-in generated by SunyBiotech for this study, crossed into N2 twice |
| IU719.1 | *rwIs33*[*hcf-1*::*gfp*::3x*flag*] *set-26(tm2467)* | Generated by genetic cross in this study |
| IU718.1 | *hcf-1(pk924) rwIs27*[*set-26*::*ha*] | Generated by genetic cross in this study |
| IU724.1 | *set-26(tm2467)*; *syb5185*[*hda-1::gfp::ha*] | Generated by genetic cross in this study |
| IU725.1 | *hcf-1(pk924)*; *syb5185*[*hda-1::gfp::ha*] | Generated by genetic cross in this study |
| IU720.1 | *glp-1(e2141)*; *rwIs33*[*hcf-1*::*gfp*::3x*flag*] | Generated by genetic cross in this study |
| IU721 | *glp-1(e2141);* *rwIs27*[*set-26*::*ha*] | Generated by genetic cross in this study |
| IU726.1 | *glp-1(e2141)*; *syb5185*[*hda-1::gfp::ha*] | Generated by genetic cross in this study |
| IU722.1 | *glp-1(e2141);rwIs33*[*hcf-1*::*gfp*::3x*flag*] *set-26(tm2467)* | Generated by genetic cross in this study |

**Supplementary Table 2. Primer sequences**

| **Target** | **Purpose** | **F Primer (5’-3’)** | **R Primer (5’-3’)** |
| --- | --- | --- | --- |
| *hda-1* | qPCR | GCACGTCATGAAGCCACATC | CAGGGAATGGGCGGAAAATC |
| *ama-1* | qPCR housekeeping | CCTACGATGTATCGAGGCAAA | CCTCCCTCCGGTGTAATAATG |
| *hda-1* | PCR amplification for verifying *syb5185* | CGTGTCGCCTACTACTATGA | TTGAGCTTCTCCCTTCCTTC |
| *hda-1* | Sanger sequencing for verifying *syb5185* | GACCAAACTACCGACTTCAC |  |
| *hda-1* | Sanger sequencing for verifying *syb5185* | GTGTTCAGGATCTGACTGTG |  |
| *set-26* | PCR amplification for verifying *rwIs27* | CGACAGTTGATGCTCCAGCTCCA | GGGGCACAAATCGAGATAGAAAGAGATGAT |
| *set-26* | Sanger sequencing for verifying *rwIs27* | CGACAGTTGATGCTCCAGCTCCA |  |
| *hcf-1* | PCR amplification of N2 *hcf-1* DNA fragments for repair template construction (upstream of stop codon) for construction of *rwIs33* | ACGTTGTAAAACGACGGCCAGTCGCCGGCAAAATCTTGAACAAAAACCACCG | CATCGATGCTCCTGAGGCTCCCGATGCTCCCTGATGATCGAAACGAGCTCTC |
| *hcf-1* | PCR amplification of N2 *hcf-1* DNA fragments for repair template construction (including and downstream of stop codon) for construction of *rwIs33* | CGTGATTACAAGGATGACGATGACAAGAGATAAACCATGGGATGGACTGATCGTTTTC | GGAAACAGCTATGACCATGTTATCGATTTCAGAGCAACCAAATCAAAGCGGA |
| *hcf-1* | Construction of gRNA for construction of *rwIs33* | ATCATCAGTAAACCATGGGAGTTTTAGAGCTAGAAATAGCAAGT |  |
| *hcf-1* | Construction of gRNA for construction of *rwIs33* | TTTCGATCATCAGTAAACCAGTTTTAGAGCTAGAAATAGCAAGT |  |
| *hcf-1* | Construction of gRNA for construction of *rwIs33* | TTCGATCATCAGTAAACCATGTTTTAGAGCTAGAAATAGCAAGT |  |
| *hcf-1* | PCR amplification for verifying *rwIs33* | ACAATCAAACCTGGGAACGGC | AGAACCTAGCTAGAGAAGAGCTCA |
| *hcf-1* | Sanger sequencing for verifying *rwIs33* | ATTGGGACAACTCCAGTGAA |  |
| *hcf-1* | Sanger sequencing for verifying *rwIs33* | GGAAGAAGAAACATCATAAACGGT |  |
| *hcf-1* | Sanger sequencing for verifying *rwIs33* | TAGATGGCTGCAGGATCAGC |  |
| *hcf-1* | Sanger sequencing for verifying *rwIs33* | TTCACTGGAGTTGTCCCAAT |  |
| *hcf-1* | Sanger sequencing for verifying *rwIs33* | AGACGATGACGATAAGCGTGAC |  |
| *hcf-1* | Sanger sequencing for verifying *rwIs33* | GGAGAACTTGTGTCCGTTGACGT |  |
| *hda-1* | *hda-1 3’ UTR* RNAi construct design | TCTGGGCCCCGAGTCGTCGAATGCAGCAAA | TCTGAGCTCTGGAAATCGTTGAAGGGGGT |

**Supplementary Table 3. Antibodies**

| **Target** | **Produced in** | **Purpose** | **Company** | **Product #** |
| --- | --- | --- | --- | --- |
| HA Tag (C29F4) | Rabbit | CUT&RUN; Immunoblot | Cell Signaling | 3724 |
| FLAG (DYKDDDDK) Tag (FG4R) | Mouse | CUT&RUN; Immunoblot | ThermoFisher | MA1-91878 |
| Histone H3 | Rabbit | CUT&RUN; Immunoblot | Abcam | ab1791 |
| Mouse IgG | Rabbit | CUT&RUN secondary (for FLAG primary) | Abcam | ab46540 |
| Rabbit IgG | Goat | Immunoblot secondary fluorescent (for HA and H3 primary) | Li-cor | 926-32211 |
| Mouse IgG | Goat | Immunoblot secondary fluorescent (for FLAG primary) | Li-cor | 926-68070 |
